# Microbial communities of the Mediterranean rocky coast: ecology and biotechnological potential

**DOI:** 10.1101/428243

**Authors:** Kristie Tanner, Esther Molina-Menor, Àngela Vidal-Verdú, Juli Peretó, Manuel Porcar

## Abstract

Microbial communities from harsh environments hold great promise as sources of biotechnologically-relevant strains. In the present work, we have deeply characterized the microorganisms from three different rocky locations of the Mediterranean coast, an environment characterised by being subjected to harsh conditions such as high levels of irradiation and large temperature and salinity fluctuations. Through culture-dependent and culture-independent techniques, we have retrieved a complete view of the ecology and functional aspects of these communities and assessed the biotechnological potential of the cultivable microorganisms. A culture-independent approach through high-throughput 16S rRNA amplicon sequencing revealed that all three locations display very similar microbial communities, suggesting that there is a stable community associated to the sampled region, with *Stanieria cyanosphaera, Rubrobacter* sp. and the families Flammeovirgaceae, Phyllobacteriaceae, Rhodobacteraceae and Trueperaceae being the most abundant taxa. Furthermore, shotgun metagenomic sequencing results were in concordance with the high-thoughput 16S rRNA, and allowed a description of the eukaryotic and archaeal members of the community, which were abundant in Ascomycota and halotolerant archaea, respectively. The culture-dependent approach yielded a collection of 100 isolates (mainly pigmented), out of which 12 displayed high antioxidant activities, as proved with two *in vitro* (hydrogen peroxide and DPPH) and an *in vivo* (model organism *C. elegans*) assays.

## Introduction

Microbial communities that inhabit harsh and extremophilic environments have been reported to be sources of biotechnologically-relevant bacteria and therefore hold great promise for the biotechnological industry [1]. For example, extremophilic microorganisms can yield enzymes such as lipases and esterases that can be used under a wide range of harsh conditions and have relevant applications in the food, detergent and biofuel industries [2]. There are many other examples of biotechnologically-relevant microorganisms from extreme environments, including the well-known *Thermus aquaticus*, that produces the widely-used *Taq* polymerase, or the hyperthermophilic biofuel-producing archaea that live in deep-sea hydrothermal vents [3, 4].

The present study focuses on the microorganisms that inhabit the rocky areas of the supralittoral zone (the area above the tide line that is subjected regularly to splash, but is not permanently underwater) of the Mediterranean coast, a harsh environment that combines exposure to radiation with other extreme conditions such as temperature fluctuations, dessication and high salinity. Although there are previous studies regarding microbial communities that inhabit the supralittoral zone and some authors have worked on the improvement of enriched media to increase the cultivability of these microorganisms [5–11], there is not and in-depth characterization of both their ecology and biotechnological potential at the same time. The extreme conditions those communities have to deal with as a result of living in the interphase between the aquatic and terrestrial worlds make them a good target for the search of biotechnologically-relevant microorganisms. For example, surface-associated microbial communities that are highly exposed to solar radiation are, on many occasions, rich in microorganisms that produce pigments, particularly, carotenoids [12–15]. These pigments play a key role in radiation tolerance [16, 17], and they are very valuable for the food, pharmacological and cosmetic industries as colorants, antioxidants and protectors against solar radiation, respectively [18]. We hypothesize that rough conditions of the supratidal zone are associated with a high diversity of biotechnologically relevant microbial taxa.

Recent advances in culturing and sequencing techniques have enabled the study of microbial communities associated to a wide range of environments, thus facilitating the discovery of new biotechnologically-relevant bacteria [15]. We present here a characterization of the bacterial communities of the sun-exposed intertidal/splash zone in the Mediterranean coast. We have compared three different coastal locations of the Mediterranean West coast and combined culturing techniques and high throughput sequencing data (16S rRNA amplicon and metagenomic sequencing) in order to shed light on the taxonomic composition of these communities, and also isolated cultivable strains with potential applications as antioxidants.

## Materials and Methods

### Sampling

Samples were collected from three different locations on the Mediterranean Western coast, in Eastern Spain: Vinaròs (Castellón), Cullera (Valencia) and Dénia (Alicante). Three samples of dark-stained rock, at least two meters apart from each other and considered as biological replicates, were collected from the supralittoral (splash) zone of each location by scraping the surface with a sterile blade. Samples of the adjacent marine water were also taken, and both types of samples (scrapped rock and sea water samples) were stored in Falcon tubes in 15% glycerol and transported to the laboratory on ice, and then stored at -20°C until required.

### High-throughput rRNA and metagenomic sequencing

Total DNA was isolated from the samples with the PowerSoil DNA Isolation kit (MO BIO laboratories, Carlsbad, CA, US) following the manufacturer’s instructions. The quantity and quality of the isolated DNA was assessed using a Nanodrop-100 Spectrophotometer (Thermo Scientific, Wilmington, DE) and purified DNA samples were sequenced by Life Sequencing SL (València, Spain) for sequencing. On one hand, the hypervariable V3-V4 regions of the 16S rRNA gene was amplified based on the protocol established by Klindworth *et al*. [19] and sequenced on the high-throughput NextSeq 500 (Illumina) platform. On the other hand, shotgun metagenomic sequencing was performed on the MiSeq Illumina platform, with paired-end sequences and reads of 150 base pairs (bp).

### Isolation and identification of bacterial strains

Three different growth mediums were used for this study: Lysogenic Broth (LB, composition in g/L: 10 tryptone, 10 NaCl, 5.0 yeast extract, 15 agar); Reasoner’s 2A agar (R2A, composition in g/L: peptone 0.5, casaminoacids 0.5, yeast extract 0.5, dextrose 0.5, soluble starch 0.5, K_2_HPO_4_ 0.3, MgSO_4_ 0.05, sodium pyruvate 0.3, 15 agar); and Marine Agar (composition in g/L: peptone 5.0, yeast extract 1.0, ferric citrate 0.1, NaCl 19.45, MgCl_2_ 5.9, Na_2_SO_4_ 3.24, CaCl_2_ 1.8, KCl 0.55, NaHCO_3_ 0.16, KBr 0.08, SrCl2 0.034, H3BO_3_ 0.022, Na_4_O_4_Si 0.004, NaF 0.024, NH4NO3 0.0016, Na_2_HPO_4_ 0.008, 15 agar). The scraped rock samples were homogenised in the Falcon tube by vigorously mixing with a vortex dilutions were cultured on the different media and incubated at room temperature for seven days. After one week of incubation, individual colonies were selected based on colony pigmentation and isolated by independent re-streaking on fresh medium. Pure cultures were then cryo-preserved at -80°C in 20% glycerol (vol:vol) until required.

Colony PCR and, were needed, DNA extracts of each of the isolated strains were used to identify them through 16S rRNA sequencing using universal primers 28F (5’-GAG TTT GAT CNT GGC TCA G-3’) and 519R (5’-GTN TTA CNG CGG CKG CTG-3’). Colony PCR cycle was performed (30 cycles of 30 s at 95°C, 30 s at 54°C, 30 s at 72°C, followed by ten minutes at 72°C and a then hold at 4°C), and included an additional initial step of incubation at 95°C for five minutes to lyse cells. The DNA extraction was done following the Latorre *et al*. (1986) protocol [20]. A PCR with a different reverse primer (1389R: 58-ACG GGC GGT GTG TAC AAG-38) was also performed for the isolates that could not be identified in any of the previous cases. Amplifications were verified by electrophoresis in a 0.8% agarose gel and then amplicons were precipitated overnight in isopropanol 1:1 (vol:vol) and potassium acetate 1:10 (vol:vol) (3 M, pH 5). DNA pellets were washed with 70% ethanol and resuspended in 30 μL Milli-Q water. BigDye^®^ Terminator v3.1 Cycle Sequencing Kit (Applied Biosystems, Carlsbad, CA, USA) was used to tag amplicons, which were sequenced with the Sanger method by the Sequencing Service (SCSIE) of the University of Valencia (Spain). The sequences were manually edited with Pregap4 (Staden Package, 2002) to eliminate low-quality base calls, and final sequences were compared by NCBI BLAST tool to nucleotide databases.

### Antioxidant activity

#### Hydrogen peroxide assay

The collection of isolates was initially screened for antioxidant activity by applying oxidative stress to the isolated colonies through the addition of hydrogen peroxide (H_2_O_2_) to the growth medium. In order to do so, all the isolates were grown on solid media for four days or until reaching enough biomass. Then, the optical density at 600 nm (OD_600_) was measured, adjusted to a value of 1, and serial dilutions were prepared up to seven times fold. Two microliters of each dilution were placed on LB or Marine Agar, to which 1 mM H_2_O_2_ had been previously added. The plates were incubated at room temperature and in the dark to avoid degradation of the H_2_O_2_, and results were recorded after two, four and six days. Two strains were used as controls for the assay: *Planomicrobium glaciei* was used as an internal positive control for antioxidant activity and *Escherichia coli* JM109 as a negative control. *P. glaciei* is a pigmented microorganism whose antioxidant activity was previously assessed *in vivo* in the lab using a *Caenorhabditis elegans* model (data not shown).

#### DPPH assay

A second assay using 2,2-diphenyl-1-picrylhydrazyl (DPPH) was performed to dismiss false positives in the H_2_O_2_ assay due to catalase activity and to confirm the antioxidant activity of the selected strains (strains with the best antioxidant activity according to the previous assay). To perform this assay, pigments were extracted from the isolates based on the protocols described by Brand-Williams *et al*. (1995), von Gadow *et al*. (1997) and Su *et al*. (2015) [21–23], with the modifications suggested by Sharma & Bhat [24]. Briefly, the isolates were grown overnight in liquid LB medium and OD_600_ was measured and normalised at a value of 1.2 for all the isolates. The cells were then harvested by centrifugation at 13,000 rpm for 3.5 min, and the pellets were resuspended in 500 μL of methanol and vigorously vortexed and sonicated for five minutes (Ultrasonic bath XUBA1, Grant Instruments, Royston, United Kingdom). The supernatant was collected by centrifugation at 13,000 rpm for three minutes, stored in a new Eppendorf tube and kept in the dark until the assay was performed. The extraction was performed twice or until a colourless pellet was obtained.

For the DPPH assay, 600 μL of the extract in methanol were mixed with 400 μL of DPPH solution (50 μM in methanol) and incubated for 30 minutes in the dark. Methanol was used as a blank and the negative control sample consisted of DPPH mixed with methanol. Absorbance was measured at 517 nm (Ultrospec 200 UV/V Visible Spectrophotometer, Pharmacia Biotech, Piscataway Township, NJ, US).

To select a suitable positive control, a standard curve with ascorbic acid (vitamin C) was performed at 10, 5, 1, 0.5, 0.1, 0.05 and 0.01 μg/mL concentrations in methanol. The detection threshold was established at 0.5 μg/mL of vitamin C, as lower concentrations of vitamin C did not change DPPH absorbance (data not shown).

DPPH scavenging ability was quantified by measuring the decrease in the absorbance of this compound at 517 nm, and the percentage of scavenged DPPH was calculated using the following formula:

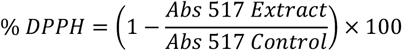

*In vivo* oxidative stress assays with *C. elegans*

The two best isolates in terms of antioxidant activity, as deduced by the results from the previous assays (CR22 and CR24) were used in an *in vivo* test with the nematode *C. elegans* as a model organism. Wild-type *C. elegans* strain N2 (Bristol, United Kingdom) was routinely propagated at 20°C on Nematode Growth Medium (NGM) plates supplemented with *E. coli* strain OP50 as the regular food source.

Nematodes were synchronized by isolating eggs from gravid adults at 20°C. Synchronization was performed on NGM plates with the different treatments: *E. coli* OP50 was supplied as a negative control, *E. coli* OP50 plus vitamin C (vitC) at 10 μg/mL as a positive control, and *E. coli* OP50 plus one of the selected isolates was used in order to test antioxidant properties of the bacteria. Duplicates were performed for every condition. Selected bacterial strains were grown overnight in liquid LB medium at 28°C and 180 rpm. Then, OD_600_ was adjusted to 30 and 50 μL of the bacterial suspension was added to the plates.

The synchronized worms were incubated for three days on the previously described plates, until reaching young adult stage. Then, young adult worms were selected for each treatment (n=50) and incubated at 20°C on the corresponding treatment, until reaching 5-day adult stage. The selected worms were then transferred to plates containing basal medium supplemented with 2 mM H_2_O_2_ and incubated for 5 hours at 20°C. After incubation, survival rates for each condition (negative control, positive control and bacteria-fed worms) were recorded by manually counting the number of living vs. dead worms.

## Results

### High-throughput 16S rRNA analysis

High throughput 16S rRNA sequencing of the samples revealed that the taxonomic composition was very similar not only within the three replicates from each location but also between samples from different locations (Fig. 1A). According to the data represented in the Krona plots (a Krona plot corresponding to one of the samples from Dénia can be seen in Fig. 1B), *Stanieria cyanosphaera* was the most abundant species in all the three locations, with its highest percentages in Cullera (38.6%) and Dénia (36%), followed by Vinaròs (28.7%). Apart from Cyanobacteria (*S. cyanosphaera*), the other dominant groups in the samples were Proteobacteria, Actinobacteria and Bacteroidetes, although their abundance varied between locations. The samples from Vinaròs were composed of 23% of Proteobacteria, 31% of Bacteroidetes and 4% Actinobacteria; whereas in Cullera and Dénia, Proteobacteria was the dominant group (with 17% and 21%, respectively), followed by Bacteroidetes (17.67% in Dénia and 13.3% in Cullera), and Actinobacteria (10.3% in Cullera and 13.6% in Dénia).

**Figure 1.**
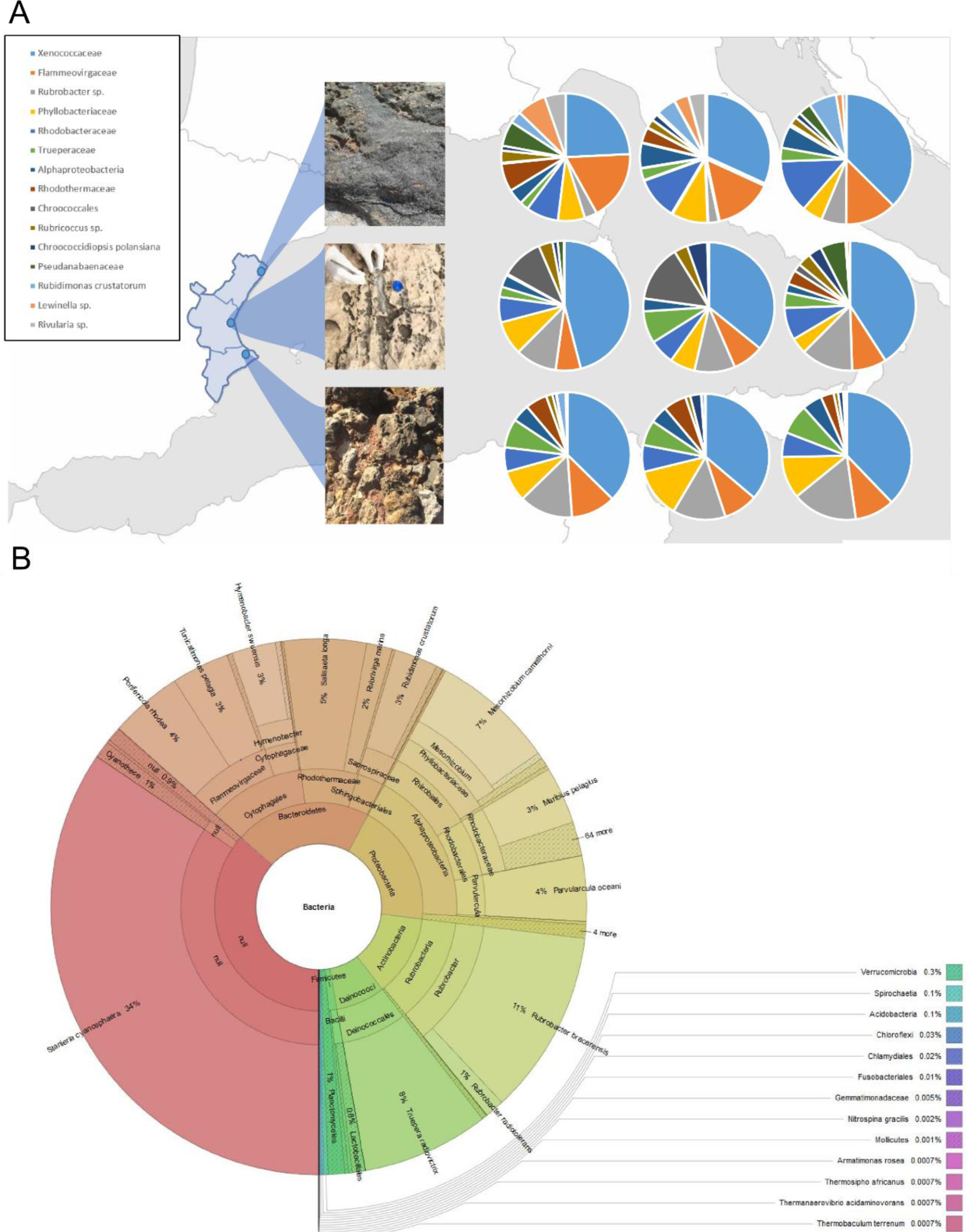
(**A**) Main taxonomic groups in samples obtained from three different locations of the Mediterranean coast (from north to south: Vinaròs, Cullera and Dénia) as obtained through high-throughput 16S rRNA amplicon sequencing. (**B**) Krona plot representing the bacterial population in one of the samples obtained from Dénia through high-throughput 16S rRNA sequencing.

The second most abundant taxon was the Flameovirgaceae family, represented by the species *Porifericola rhodea* and *Tunicatimonas pelagia*, which appeared with similar frequencies in Dénia and Cullera (4 and 4.67% in Dénia and 3.67 and 3% in Cullera, respectively), and 9% *(P. rhodea)* and 2.67% *(T. pelagia)* in Vinaròs. The genus *Rubrobacter* was mainly represented by *Rubrobacter bracarensis* (12.7, 7.3 and 2.33% in Dénia, Cullera and Vinaròs, respectively), and *Rubrobacter radiotolerans* (2.33, 1 and 0.33% in Cullera, Vinaròs and Dénia, respectively), but other species were also found in very low frequencies, such as *Rubrobacter taiwanensis* and *Rubrobacter naiadicus*. Interestingly, the radiation-resistant thermophilic bacteria *Truepera radiovictrix* was also detected in all the sampled locations, with abundances ranging from 2.67% in Vinaròs to 7.33% in Dénia.

**Figure 2.**
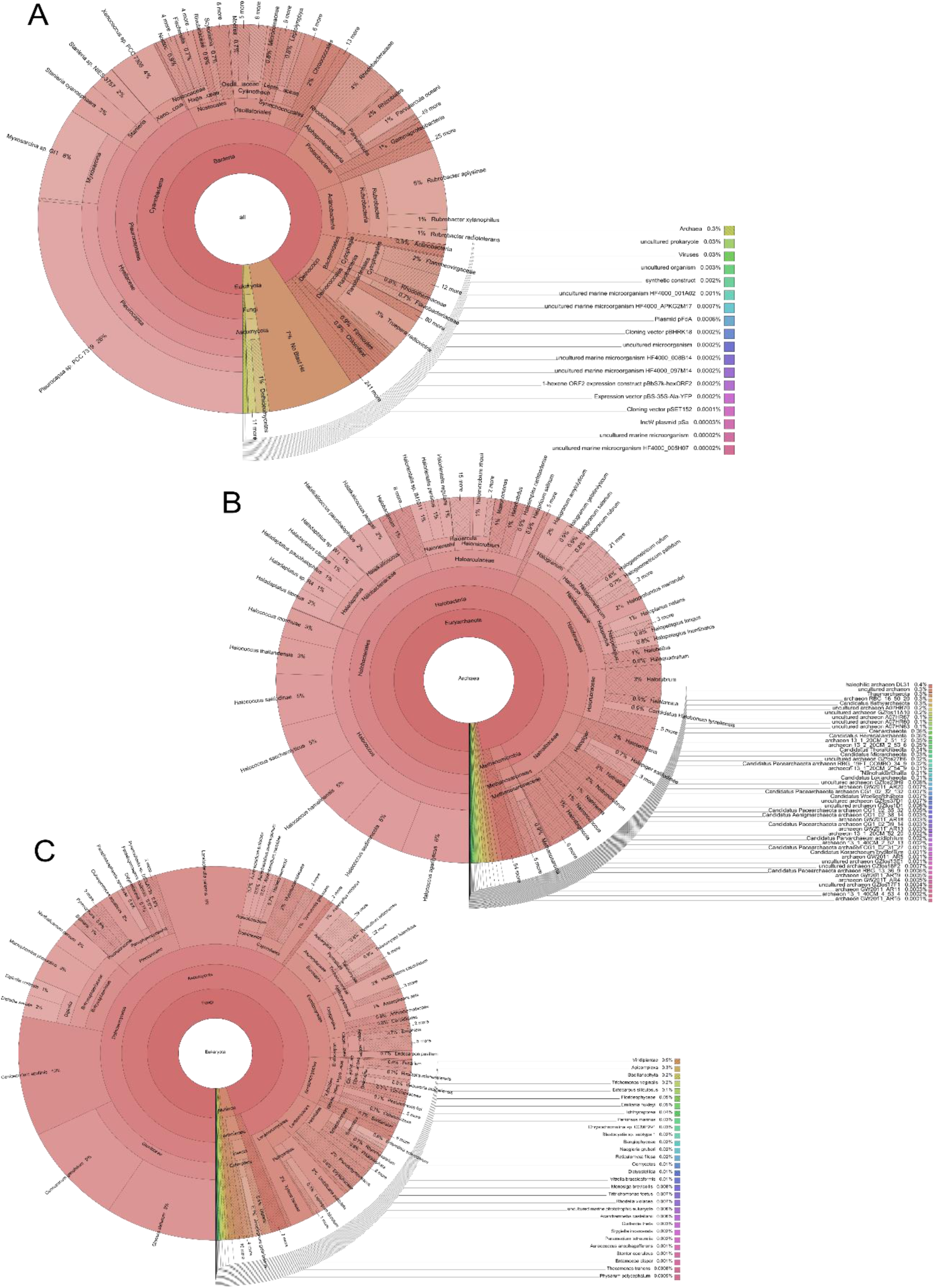
Main bacterial (**A**), archaeal (**B**) and eukaryotic (**C**) groups identified in the sample obtained from Dènia and analysed through metagenomic sequencing.

### Shotgun metagenomic analysis

Once again, all three replicates proved very similar in terms of the taxonomic profile (data not shown). Although the most abundant bacterial phyla were the same as the ones observed with high-throughput 16S rRNA sequencing, the abundance of these varied slightly, with Cyanobacteria being the most abundant in all cases, followed by Proteobacteria, Actinobacteria and Bacteroidetes, and with a lower amount of Deinococcus-Thermus (Figure 2A). Furthermore, the abundance of *S. cyanosphaera* was lower than the abundance obtained when analysed through high-throughput 16S rRNA sequencing, and metagenomic sequencing revealed other abundant taxa in the Cyanobacteria phyla, including the genera *Pleurocapsa*, *Myxosarcina* and *Xenococcus*. The overall taxonomic profiles proved very similar with both types of sequencing techniques.

Archaeal and eukaryotic communities proved very diverse, with a high number of salt-adapted microorganisms in the former and a large fraction of Ascomycota in the latter (**Figure 2B and 2C**). Salt-adapted archaea included members of *Halococcus*, Halobacteriaceae *(Haladaptatus* and *Halalkalicoccus)*, Haloarculaceae, Haloferaceae, Halorubraceae and Natrialbaceae, and methanogenic archaea were also detected (specifically members of the Methanosarcinaceae family (**Figure 2B**)). Among the diversity of Ascomycota, the most abundant taxa were *Glonium stellatum*, *Cenococcum geophilum*, *Coniosporium apollinis* and *Lepidopterella palustris* (**Figure 2C**).

### Strain collection and identification

Culturing the samples on LB and Marine Agar medium yielded a large diversity of colonies in terms of colour, shape and morphology. The majority of the isolates could grow on both LB and Marine Agar, while few of them could on R2A. A total of 100 strains were isolated and named with the following code, after the location (C: Cullera, D: Dénia, V: Vinaròs), and the origin (M: Marine water, R: Rock surface). There was no significant fungal growth in any of the samples. The colonies observed on Marine Agar displayed the widest range of colours (wine-red, red, pink, and orange, among others) in comparison with the ones observed on LB media, which were mainly yellowish and cream-coloured. Due to the known relation between the presence of pigments and antioxidant power, the main criterion for colony selection was the colour [25].

A collection of 100 isolates in pure culture was established and cryo-preserved in 20% glycerol. A total of 34 isolates were identified through colony PCR and 16S rRNA Sanger sequencing. Although an initial step of incubation at 100°C was added to the PCR protocol of the isolates whose amplification had failed, some remained non-identified and therefore their total DNA was extracted to repeat the PCR. Finally, 56 of the isolates still remained unidentified. Among the identified isolates, there were many *Bacillus* spp. *(B. alkalitelluris, B. amyloliquefaciens*, *B. aquimaris*, *B. marisflavi*, *B. mojavensis*, *B. salsus* and *B. subtilis*), as well as other species such as *Halobacillus* spp., *Micrococcus antarcticus, Micrococcus luteus*, *Staphylococcus* spp., *Vibrio corallilyticus*, *Vibrio tubiashii* and *Virgibacillus halodenitrificans*.

### Antioxidant activity

In order to select and establish a collection of isolates with antioxidant properties, a high-throughput screening of the 100 isolates was performed by growing them on solid media containing H_2_O_2_. *P. glaciei* and *E. coli* JM109 were used as positive and negative controls, respectively. Strain JM109, with no known reports of antioxidant effect, exhibited a weak growth in the first (OD_600_ 1) and, sometimes, second dilution (OD_600_ 10^−1^). This lead us to the criterion to consider positive antioxidant activity those strains able to grow on H_2_O_2_-containing plates at least up to three-fold diluted (OD_600_ 10^−2^). A total of 12 isolates were selected (Table 1) based on their ability to grow on 1 mM H_2_O_2_ plates as described above.

DPPH-based assays are widely used to detect and quantify the antioxidant power of plants or bacterial extracts. These assays are based on the decrease of DPPH absorbance at 517 nm in presence of antioxidant factors. The oxidative stress-resistant isolates selected from the H_2_O_2_ assay (shown in **Table 1**) were further tested using this method. CR17, CR21 and CR57 could not be tested due to poor growth in liquid culture that made it impossible to obtain a concentrated extract. 16S rRNA sequences were compared using NCBI BLAST tool. Isolates CR10-VR2 and CR22-CR28 were 100% identical in their 16S rRNA sequence, and therefore only one of each was selected for further assays.

The test resulted in a general decrease in absorbance in all the samples, suggesting that the extracts were able to scavenge the DPPH. The isolates that proved to be more effective as antioxidants were CR22, CR24 and CR28, with values of scavenged DPPH over 30% (Figure 3A). DR12 displayed low DPPH scavenging values possibly due to failure of the pigment extraction. Surprisingly, the control samples *P. glaciei* and JM109 did not display the expected effect. A set of both theoretically positive and negative controls for antioxidant activity were tested through DPPH for a deeper undersntanding of the results described above, and showed similar results (**Figure 4**)

**Table 1.** List of selected isolates, percentage of identity with the closest Blast, identification and results obtained through H_2_O_2_ assay (number of dilutions able to grow on H_2_O_2_ containing medium).

| Sample | Taxonomic Identification | H<sub>2</sub>O<sub>2</sub> Assay |
| --- | --- | --- |
| CR10 | *Micrococcus luteus* (99%) | 3 |
| CR17 | *Virgibacillus halodenitrificans* (99%) | 7 |
| CR21 | Non-identified | 4 |
| CR22 | *Virgibacillus halodenitrificans* (99%) | 4 |
| CR24 | *Halobacillus* sp. (99%) | 6 |
| CR28 | *Virgibacillus halodenitrificans* (99%) | 6 |
| CR37 | *Bacillus* sp. (99%) or *Bacillus aquimaris* (99%) | 4 |
| CR44 | Non-identified | 3 |
| CR67 | *Bacillus* sp. (99%) or *Bacillus aquimaris* (99%) | 4 |
| DM10 | Non-identified | 3 |
| DR12 | Non-identified | 3 |
| VR1 | *Vibrio corallilyticus* (99%) or *Vibrio tubiashii* (99%) | 6 |
| VR2 | *Micrococcus luteus* (99%) | 3 |
| Positive control | *P. glaciei* | 8 |
| Negative control | *E. coli* JM109 | 1 |

The two strains with best results from the *in vitro* assays (CR22 and CR24) were selected for further *in vivo* antioxidant assays in the model organism *C. elegans*, and both strains displayed an important antioxidant activity (**Figure 3B**). Worms that were subjected to oxidative stress after being treated with isolates CR22 and CR24 displayed survival rates higher than the untreated worms and similar to those observed in the worms treated with vitamin C (survival rates of around 55–65%), a well-known antioxidant used as a positive control.

## Discussion

We report here, for the first time, and by using culture-dependent and independent (NGS) techniques the microbiomes of the rocky-coastal surface of three regions on the Mediterranean western coast. The three sampled locations displayed rather similar taxonomic profiles, suggesting that the microbial composition of the Mediterranean supratidal zone, at least in eastern Spain, is very stable. The studied communities were particularly dominated by bacterial strains previously described as thermophilic, halotolerant or radioresistant, such as the species *S. cyanosphaera* or the genus *Rubrobacter* [26, 27], and pigmented isolates, as is the case of the most abundant species within the Flameovirgaceae family, *T. pelagia* and *P. rhodea* [28, 29].

Interestingly, *T. radiovictrix*, characterised by an optimum growth temperature at 50<u>^°^</u>C and an extreme resistance to ionizing radiation, was first recovered from a hot spring in a geothermal area close to the Azores [30, 31]. Moreover, the *Truepera* genus has been previously found in Lake Lucero Playa (New Mexico, USA), a particularly hostile environment as the lake dries periodically [32]. This is, to the best of our knowledge, the first report of sea-inhabiting *Truepera* spp. in a non-thermal environment and it is tempting to hypothesize that *Truepera* spp. might have a similar ecological niche than Deinococcus, but in saline environments, as a consequence of both its radiation resistance and halotolerance [30].

Shotgun metagenomic analysis confirmed the similarity between the communities of the three sampled locations, as discussed above from the high-throughput 16S rRNA results, particularly at higher taxonomic levels (i.e. family level) were similar. Nevertheless, the results at lower taxonomic levels varied considerably among sequencing techniques. One of the largest differences at the species level was observed within the Cyanobacterial group. In particular, high-throughput 16S rRNA revealed a large abundance of *S. cyanosphaera*, whereas shotgun metagenomic sequencing revealed a more diverse population including members of *Pleurocapsa*, *Myxosarcina* and *Xenococcus*, as previously reported for marine environments [33–36].

**Figure 3.**
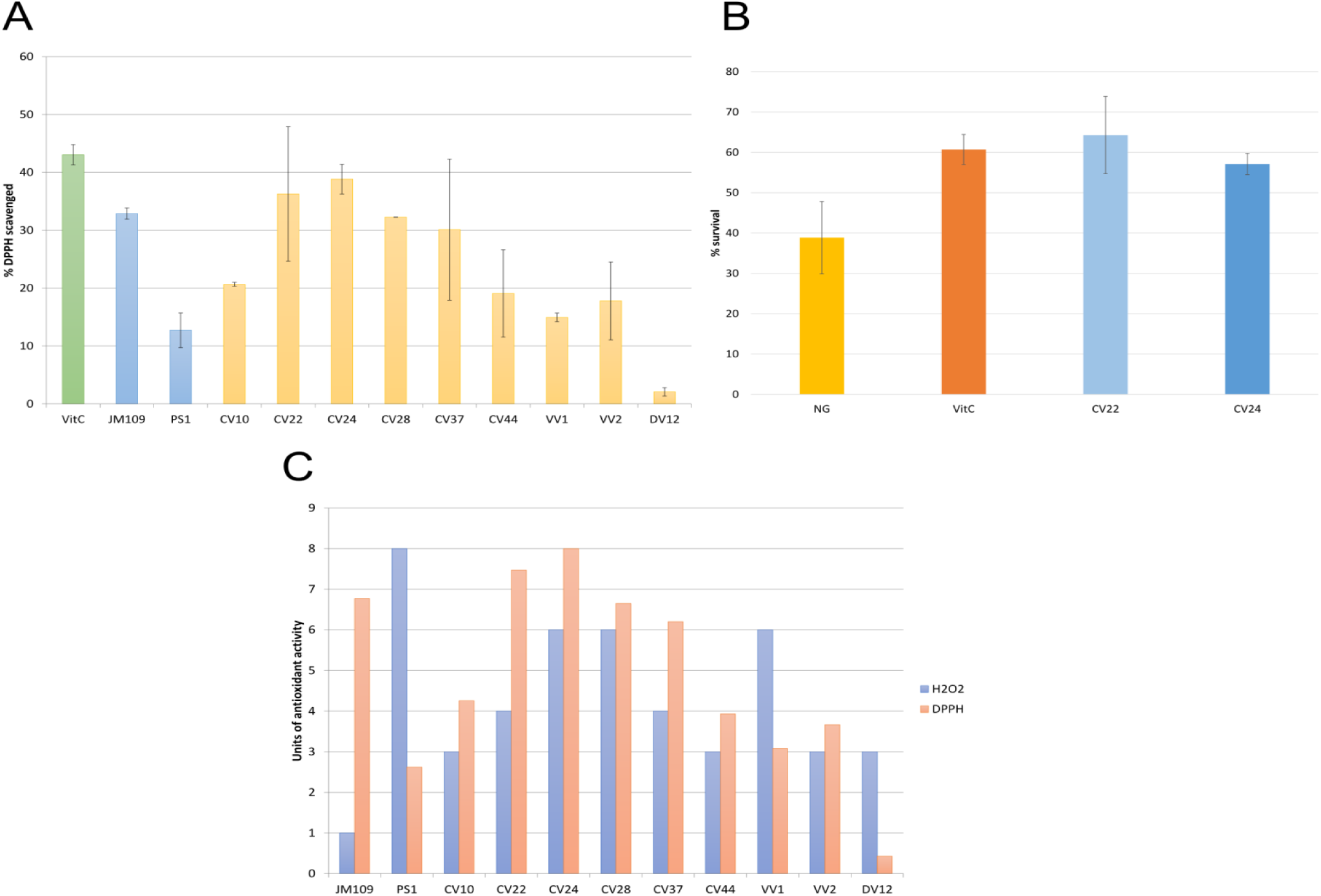
(**A**) DPPH assay. Absorbance was measured at 517 nm after 30 minutes of incubation with DPPH 50 μM. DPPH scavenged percentage is represented in Y-axis. VitC, vitamin C (0.5 μg/mL solution); (B) Antioxidant activity in vivo (using the model organism C. elegans). Y-axis indicates percentage of survival of worms after five hours of incubation under oxidative stress (H_2_O_2_). Worms were treated with either a control diet (NG), a diet supplemented with the known antioxidant vitamin C as a positive control (VitC) or a diet supplemented with the selected strains (CR22 and CR24). (C) Comparative analysis of H_2_O_2_ and DPPH assay. Values in Y-axis are normalised with respect to the highest value obtained in both assays..

The large abundance and diversity of salt-adapted archaea in the sampled marine environments could be explained by the adaptation adaptative characteristics developed by these microorganisms, which allow them to survive to desiccation, starvation and radiation through several different strategies, including the formation of halomucin or the survival in fluid inclusions of halite [37]. Furthermore, previous reports described the salt adaptation mechanisms of *Methanosarcina mazei*, which includes accumulation of solutes and export of Na^+^ [38].

The eukaryotic fraction of the samples was mainly composed by Ascomycota, such as *Glonium stellatum*. The genus *Glonium* includes saprophytic Dothideomycetes that produce darkly-pigmented apothecia, which could explain the dark colour observed in the sampled rocks [39]. Other species detected in the samples included: *Cenococcum geophilum*, an ectomycorrhizal fungus previously described in coastal forest soils [40] and previously demonstrated to grow at up to 100 mM of NaCl [41]; *Coniosporium apollinis*, a rock-inhabiting fungi previously isolated from the Mediterranean basin [42]; and *Lepidopterella palustris*, typically a freshwater fungi [43], with this being, to the best of our knowledge, the first description of this species in a salt water habitat.

Taken together, the results obtained from both high-throughput 16S rRNA and metagenomic sequencing suggest that the sampled communities are composed of a diverse array of fungi (mainly belonging to the phylum Ascomycota), cyanobacteria (mainly *S. cyanosphaera* and *Pleurocapsa* spp., but also *Myxosarcina* spp. and *Xenococcus* spp.) and salt-adapted archaea, which remain stable among the three different sampled locations.

**Figure 4.**
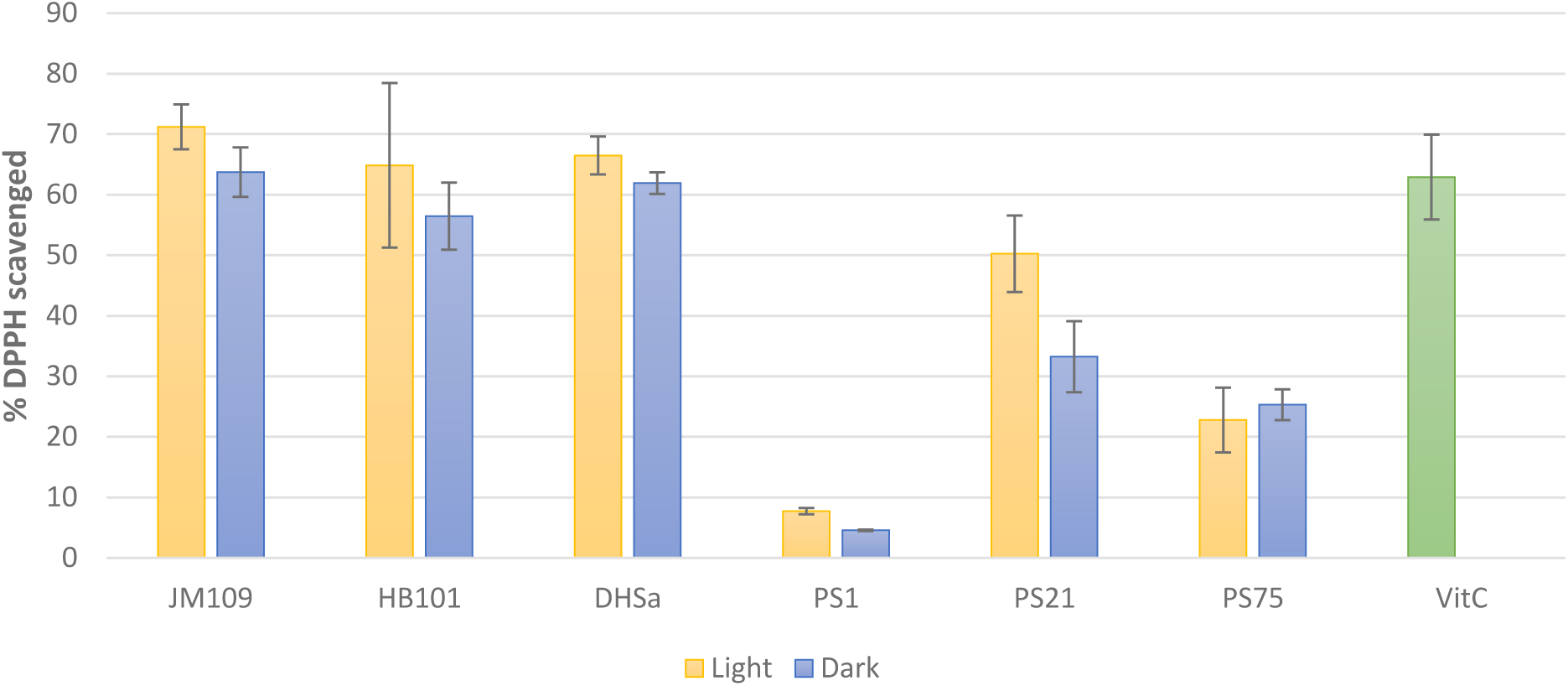
DPPH assay with positive and negative controls. Absorbance was measured at 517 nm after 30 minutes of incubation with DPPH 50 μM. DPPH scavenged percentage is represented in Y-axis. VitC, Vitamin C, 0.5 μg/mL solution. Light and dark conditions of growth are represented.

From the collection of cultured microorganisms, a total of 12 isolates were selected as positive for antioxidant activity through an oxidative stress assay performed with H_2_O_2_. Of these, of *M. luteus* has been reported to encode genes related to resistance and tolerance to oxidative stress (superoxide dismutase and NADP reductase) [44]. On the other hand, *Vibrio* species associated with corals, such as *Vibrio corallilyticus*, produce enzymes that react with reactive oxygen species (ROS), not only through catalase but also through superoxide dismutase (SOD) [45]. The DPPH assay was performed to dismiss false positives through the H_2_O_2_ assay. In general, the results correlated well with the ones previously observed in the H_2_O_2_ assay. It is important to note that, although DR12 displayed low scavenging in the DPPH assay, the extraction of pigments from this isolate was sub-optimal, since the pellet remained pink-coloured after the extraction process. Surprisingly, the control samples *P. glaciei* and JM109 did not display the expected effect. On one hand, *P. glaciei* was expected to be one of the most antioxidant isolates, as its antioxidant activity was demonstrated in previous *in vivo* assays in *C. elegans* (data not shown) and in the H_2_O_2_ assay. Nevertheless, it was the worst dtrain in terms of DPPH scavenging. On the other hand, *E. coli JM109*, with no previous reports on antioxidant activity, resulted in high DPPH scavenging.

In general, though, the correlation between both methods appears to be, in general, good, as the isolates with higher survival in the presence of H_2_O_2_ also displayed higher DPPH-scavenging ability (**Figure. 3C**). Nevertheless, there were some isolates that displayed different results depending on the method, in particular VR1 and CR37. Differences in VR1 could be the result of catalase activity, which may have enhanced its growth on the H_2_O_2_-supplemented plates. On the contrary, differences between both methods for CR37 could be caused by a deficient growth in solid medium. This results highlight the limitation of screening techniques for antioxidant activities.

A collection of both positive and negative controls (in terms of theoretical antioxidant activity) were tested using both assays (H_2_O_2_ and DPPH). *P. glaciei, Paracoccus* spp. and *Bacillus megaterium* were the three control strains selected, all of them recovered from solar panels and previously tested in *C. elegans* for in vivo protection against oxidative stress (data not shown), and three different strains of *E. coli* were chosen as negative controls (JM109, BH101, DH5α). For the DPPH assay, the isolates were grown in both light and dark conditions, in order to determine whether the light had a negative impact on the production of pigments or other antioxidant factors, as it is known that many pigments, particularly carotenoids, are prone to photodegradation [46]. For the *E. coli* strains, no significant differences were observed between growth in dark and light conditions, whereas PS21 proved very sensitive to light (**Figure 4**). Moreover, the scavenging effect of the JM109 strain that had previously been detected could be reproduced and observed in the other two *E. coli* strains as well, confirming that the extracts obtained from *E. coli* contain compounds are indeed able to react with DPPH. Even though *Paracoccus* spp. and *B. megaterium* displayed better antioxidant properties than *P. glaciei*, which was again comparable to the negative control of methanol, they were apparently less effective than the *E. coli* strains, according to this assay.

Taken together, our results suggest that the western litoral rocky coast the Mediterranean sea harbours a stable microbial community that is conserved among different locations, with the cyanobacteria *S. cyanosphaera* as the majoritarian bacterial taxa, followed by members of the Flameovirgaceae family and members of the *Rubrobacter* genus, as well as eukaryotic and archaeal members, such Ascomycota and halotolerant archaea. Furthermore, *in vitro* and *in vivo* assays have revealed that this environment is a potential source of microorganisms with antioxidant activities that can be further used for a wide range of applications in the food, cosmetic and pharmacological industry.

## Acknowledgments

We thank ADM-Biopolis S.L. for granting us access to their laboratory and materials for the *C. elegans* assays. We also thank Adriel Latorre and Darwin Bioprospecting Excellence S.L. (Valencia, Spain) for his assistance with the bioinformatic analysis.

## Competing Interests

The authors declare no conflict of interest.

## Author contributions

MP conceived the work. MP, KT and EMM collected the samples. KT, EMM and AVV performed the culture-based characterization, and KT carried out the bioinformatic analysis. All authors (MP, KT, AVV, EMM and JP) analysed the results, wrote and approved the manuscript.

## Funding

Financial support from Spanish Government (grant Helios. reference: BIO2015-66960-C3-1-R co-financed by FEDER funds and Ministerio de Economía y Competitividad) is acknowledged.

## References

[1] Raddadi N, Cherif A, Daffonchio D, Neifar M and Fava F. Biotechnological applications of extremophiles, extremozymes and extremolytes. Appl Microbiol Biotechnol. 2015; 99: 7907.

[2] Fuciños P, González R, Atanes E, Sestelo AB, Pérez-Guerra N, Pastrana L, et al. Lipases and esterases from extremophiles: overview and case example of the production and purification of an esterase from Thermus thermophilus HB27. Methods. Mol Biol. 2012;861: 239–66.

[3] Chien A., Edgar DB., and Trela JM. Deoxyribonucleic acid polymerase from the extreme thermophile Thermus aquaticus. J Bacteriol. 1976;127: 1550–1557

[4] Nishimura H. and Sako Y. Purification and characterization of the oxygen-thermostable hydrogenase from the aerobic hyperthermophilic archaeon Aeropyrum camini. J Biosci Bioeng. 2009;108, 299–303.

[5] Bates, S. T., Cropsey, G. W., Caporaso, J. G., Knight, R., and Fierer, N. Bacterial Communities Associated with the Lichen Symbiosis. Applied Environ Microb. 2011;77 (4):1309–14.

[6] Cardinale M., Puglia A. M. and Grube M. Molecular analysis of lichen-associated bacterial communities. FEMS Microbiol Ecol. 2006;57:484–95

[7] González I., Ayuso-Sacido A., Anderson A. and Genilloud O. Actinomycetes isolated from lichens: evaluation of their diversity and detection of biosynthetic gene sequences. FEMS Microbiol. Ecol. 2005;54:401–15.

[8] Grube M. & Berg G. Microbial consortia of bacteria and fungi with focus on the lichen symbiosis. Fungal Biol Rev. 2009;23:72–85.

[9] Hodkinson B. P. and Lutzoni F. A microbiotic survey of lichen-associated bacteria reveals a new lineage from the Rhizobiales. Symbiosis 2010;49:163–80.

[10] Sánchez-Hidalgo, M., González, I., Díaz-Muñoz, C., Martínez, G., & Genilloud, O. Comparative Genomics and Biosynthetic Potential Analysis of Two Lichen-Isolated Amycolatopsis Strains. Front Microbiol, 2018;9:369.

[11] Biosca E.G, Flores, R., Santander, R., Díez-Gio, J.L., Barreno, E. Innovative approaches using lichen enriched media to improve isolation and culturability of lichen associated bacteria. PLOS ONE. 2016; doi.org/10.1371/journal.pone.0160328

[12] Dorado-Morales, P., Vilanova, C., Peretó, J., Codoñer, F.M., Ramón, D. and Porcar, M. A highly diverse, desert-like microbial biocenosis on solar panels in a Mediterranean city. Sci Rep-UK. 2015;6, 29235. doi: 10.1038/srep29235.

[13] Kumar, V.B.N., Kampe, B., Rösch, P. and Popp, J. Characterization of carotenoids in soil bacteria and investigation of their photo degradation by UVA radiation via resonance Raman spectroscopy. Analyst. 2015;140:4584–93.

[14] Shindo, K. and Misawa, N. New and rare carotenoids isolated from marine bacteria and their antioxidant activities. Marine drugs 2014;12:1690–8.

[15] Tanner, K., Vilanova, C., Porcar, M. Bioprospecting challenges in unusual environments. Microb Biotechnol. 2017;10(4):671–3.

[16] Tian, B. and Hua, Y. Carotenoid biosynthesis in extremophilic Deinoccocus-Thermus bacteria. Trends in Micorbiology 2010;18(11):512–20.

[17] Tanner, K., Martí, J.M., Belliure, J., Fernández-Méndez, M., Molina-Menor, E., Peretó, J., and Porcar, M. Polar solar panels: Arctic and Antarctic microbiomes display similar taxonomic profiles. Env Microbiol Rep. 2018;10(1):75–9.

[18] Sandmann, G. Carotenoids of biotechnological importance. Adv Biochem Eng Biot. 2015;148:449–67.

[19] Klindworth, A., Pruesse, E., Schweer, T., Peplies, J., Quast, C., Horn, M., et al. Evaluation of general 16S ribosomal RNA gene PCR primers for classical and next generation sequencing-based diversity studies. Nucleic Acids Res, 2013;41(1):e1.

[20] Latorre, A., Moya, A., & Ayala, F. Evolution of mitochondrial DNA in Drosophila subobscura. P Natl Acad Sci. 1986;83:8649–53.

[21] Brand-Williams, W., Cuvelier, M., & Berset, C. Use of a Free Radical Method to Evaluate Antioxidant Activity. Lebensmittel-Wissenschaft & Technologie. 1995;28:25–30.

[22] von Gadow, A., Elizabeth, J., & Hansmann, C. Comparison of the Antioxidant Activity of Aspalathin with That of Other Plant Phenols of Rooibos Tea (Aspalathus linearis), r-Tocopherol, BHT, and BHA. Journal of Agricultural and Food Chemistry. 1997;45:632–8.

[23] Su, J., Wang, T., Li, Y.-Y., Li, J., Zhang, Y., Wang, Y., et al. Antioxidant properties of wine lactic acid bacteria: Oenococcus oeni. Appl Microbiol and Biotechnol. 2015; 99: 5189–202.

[24] Sharma, O. P., and Bhat, T. K. DPPH antioxidant assay revisited. Food Chem, 2009;113:1202–5.

[25] Pawar, R., Mohandass, C., Sivaperumal, E., Sabu, E., Rajasabapathy, R., and Jagtap, T. Epiphytic marine pigmented bacteria: A prospective source of natural antioxidants. Braz J Microbiol. 2015;46 (1):29–39.

[26] Prihantini, N. B., Sjamsuridzal, W., and Yokota, A. Description of Stanieria strain of cyanobacteria isolated from hot spring in Indonesia. American Institute of Physics, AIP Conference Proceedings. 2016;1729, 020066.

[27] Jurado, V., Miller, A., Alias-Villegas, C., Laiz, L., & Saiz-Jimenez, C. Rubrobacter bracarensis sp. nov., a novel member of the genus Rubrobacter isolated from a biodeteriorated monument. Syst Appl Microbiol. 2012;35:306–9.

[28] Yoon J, Oku N, Park S, Katsuta A, Kasai H. Tunicatimonas pelagia gen. nov., sp. nov., a novel representative of the family Flammeovirgaceae isolated from a sea anemone by the differential growth screening method. Antonie Van Leeuwenhoek 2012; 101(1):133–40.

[29] Yoon J, Oku N, Park S, Kasai H, Yokota A. Porifericola rhodea gen. nov., sp. nov., a new member of the phylum Bacteroidetes isolated by the bait-streaked agar technique. Antonie Van Leeuwenhoek. 2011; 100(1):145–53.

[30] Albuquerque, L., Simoes, C., Nobre, M. F., Pino, N. M., Battista, J. R., Silva, M. T., et al. Truepera radiovictrix gen. nov., sp. nov., a new radiation resistant species and the proposal of Trueperaceae fam. nov. FEMS Microbiol Lett. 2005;247:161–9.

[31] Ivanova, N., Rohde, C., Munk, C., Nolan, M., Lucas, S., Rio, T. G., et al. Complete genome sequence of Truepera radiovictrix. Stan Genomic Sci. 2011;4:91–6.

[32] Sirisena, KA, Ramírez, ∫, Steele, A, Glamodija, M. (2018) Microbial diversity of hypersaline sediments from lake lucero playa in white sands national monument, New Mexico, USA. Microb Ecol. 76(2):404–418.

[33] Brito, A., Ramos, V., Mota, R., Lima, S., Santos, A., Vieira, J. et al. Description of new genera and species of marine Cyanobacteria from the Portuguese Atlantic coast. Mol Phylogenet Evol. 2017;111:18–34.6

[34] Yu, C.H., Lu, C.K., Su, H.M., Chiang, T.Y., Hwang, C. C., Liu, T. Chen, Y.M. Draft genome of Myxosarcina sp. strain GI1, a baeocytous cyanobacterium associated with the marine sponge Terpios hoshinota. Stand Genomic Sci. 2015;10:28.

[35] Alex, A., Vasconcelos, V., Tamagnini, P., Santos, A., Antunes, A. Unusual Symbiotic Cyanobacteria Association in the Genetically Diverse Intertidal Marine Sponge Hymeniacidon perlevis (Demospongiae, Halichondrida). PLOS ONE. 2012;7(12): e51834.

[36] Burns, B.P., Goh, F., Allen, M., Neilan, B.A. Microbial diversity of extant stromatolites in the hypersaline marine environment of Shark Bay, Australia. Environ Microbiol. 2004; 6(10):1096–101.

[37] Stan-Lotter, H. and Fendrihan S. Halophilic Archaea: Life with Desiccation, Radiation and Oligotrophy over Geological Times. Life. 2015; 5(3):1487–96.

[38] Spanheimer, R. and Müller, V. The molecular basis of salt adaptation in Methanosarcina mazei Gö1. Arch Microbiol. 2008;190(3):271–9.

[39] Spatafora, J.W., Owensby, C.A., Douhan, G.W., Boehm, E.W. and Schoch, C.L. Phylogenetic placement of the ectomycorrhizal genus Cenococcum in Gloniaceae (Dothideomycetes). Mycologia. 2012; 104(3):758–765.

[40] Matsuda, Y., Takeuchi, K., Obase, K., Ito, S. Spatial distribution and genetic structure of Cenococcum geophilum in coastal pine forests in Japan. FEMS Microbiol Ecol. 2015;91(10).

[41] Obase, K., Lee, J.K, Lee, S.K., Lee, S.Y., Chun, K.W. Variation in Sodium Chloride Resistance of Cenococcum geophilum and Suillus granulatus Isolates in Liquid Culture. Mycobiology. 2010;38(3): 225–8.

[42] Sterflinger, K., De Baere, R., de Hoog, G.S., De Wachter, R., Krumbein, W.E., Haase, G. Coniosporium perforans and C. apollinis, two new rock-inhabiting fungi isolated from marble in the Sanctuary of Delos (Cyclades, Greece). Antonie Van Leeuwenhoek. 1997;72(4): 349–363.

[43] Shearer, C.A., Raja, H.A., Miller, A.N., Nelson, P., Tanaka, K., Hirayama, K., Marvanová, L., Hyde, K.D., Zhang, Y. The molecular phylogeny of freshwater Dothideomycetes. Stud Mycol. 2009;64:145–53.

[44] Lafi, F.F., Ramírez-Prado, J_S_, Alam, I., Bajic, V.B., Hirt, H., Saad, M.M. Draft genome sequence of plan growth-promoting Micrococcus luteus strain K39 isolated grom Cyperus conglomeratus in Saudi Arabia.Genome Announc. 2017;5(4): pii: e01520–16

[45] Munn, C.B., Marchant, H.K., Moody, A.J. Defences against oxidative stress in vibrios aasociated with corals. FEMS Microbiol Lett. 2008;281(1).

[46] Boon, C.S., McClements, D.J., Weiss, J., Decker, E.A. Factors influencing the chemical stability of carotenoids in foods. Crit Rev Food Sci Nutr. 2010;50(6):515–32.

